# Intracellular replication of *Pseudomonas aeruginosa* in epithelial cells requires suppression of the caspase-4 inflammasome

**DOI:** 10.1101/2023.02.13.528260

**Authors:** Abby R Kroken, Keith A Klein, Patrick S Mitchell, Vincent Nieto, Eric J Jedel, David J Evans, Suzanne M J Fleiszig

## Abstract

Pathogenesis of *Pseudomonas aeruginosa* infections can include bacterial survival inside epithelial cells. Previously, we showed this involves multiple roles played by the type three-secretion system (T3SS), and specifically the effector ExoS. This includes ExoS-dependent inhibition of a lytic host cell response that subsequently enables intracellular replication. Here, we studied the underlying cell death response to intracellular *P. aeruginosa*, comparing wild-type to T3SS mutants varying in capacity to induce cell death and that localize to different intracellular compartments. Results showed that corneal epithelial cell death induced by intracellular *P. aeruginosa* lacking the T3SS, which remains in vacuoles, correlated with activation of NF-κB as measured by p65 relocalization and TNFα transcription and secretion. Deletion of caspase-4 through CRISPR-Cas9 mutagenesis delayed cell death caused by these intracellular T3SS mutants. Caspase-4 deletion also countered more rapid cell death caused by T3SS effector-null mutants still expressing the TSSS apparatus that traffic to the host cell cytoplasm, and in doing so rescued intracellular replication normally dependent on ExoS. While HeLa cells lacked a lytic death response to T3SS mutants, it was found to be enabled by interferon gamma treatment. Together, these results show that epithelial cells can activate the noncanonical inflammasome pathway to limit proliferation of intracellular *P. aeruginosa*, not fully dependent on bacterially-driven vacuole escape. Since ExoS inhibits the lytic response, the data implicate targeting of caspase-4, an intracellular pattern recognition receptor, as another contributor to the role of ExoS in the intracellular lifestyle of *P. aeruginosa*.

**Importance:** *Pseudomonas aeruginosa* can exhibit an intracellular lifestyle within epithelial cells in vivo and in vitro. The type three secretion system (T3SS) effector ExoS contributes via multiple mechanisms, including extending the life of invaded host cells. Here, we aimed to understand the underlying cell death inhibited by ExoS when *P. aeruginosa* is intracellular. Results showed that intracellular *P. aeruginosa* lacking T3SS effectors could elicit rapid cell lysis via the non-canonical inflammasome pathway. Caspase-4 contributed to cell lysis even when the intracellular bacteria lacked the entire T33S and were consequently unable to escape vacuoles, representing a naturally occurring subpopulation during wildtype infection. Together, the data show the caspase-4 inflammasome as an epithelial cell defense against intracellular *P. aeruginosa*, and implicate its targeting as another mechanism by which ExoS preserves the host cell replicative niche.

## Introduction

*Pseudomonas aeruginosa* is an opportunistic pathogen able to cause life- and vision-threatening infections. While often referred to as an extracellular pathogen, *P. aeruginosa* can adopt an intracellular lifestyle in various epithelial cell types, including corneal (1), bronchial (2), HeLa cells (3) and in vivo (4). The ability of *P. aeruginosa* to replicate inside a host cell depends on the type three secretion system (T3SS), with multiple roles played by the T3SS effector ExoS (5–7) encoded by only invasive strains (8). ExoS is a bifunctional protein with both RhoGAP and ADP ribosyltransferase (ADPr) activity. Broad substrate specificity of its ADPr domain (9, 10) enables ExoS to impact multiple host processes (11, 12). Demonstrated effects of ExoS include inactivation of host cell proliferative/survival signaling, disassembling the host cell cytoskeleton, freezing of intracellular membrane trafficking, and disruption of ROS generation (9, 13–16). While the molecular components of host cells targeted by ExoS to enable *P. aeruginosa* to replicate intracellularly have not yet been elucidated, its ADPr activity is required, and mechanisms shown include inhibition of autophagy (17) and evasion of lysosomes (18). Recently, we reported another role for the ADPr activity of ExoS in intracellular persistence: preservation of host cell viability (19), which was further explored here.

Eukaryotic host cells can sense and respond to intracellular pathogens (20, 21). Responses can include initiation of regulated cell death pathways (22), which may subsequently be suppressed by the bacteria to preserve their intracellular niche (23). One type of host response is pyroptosis, a form of inflammatory lytic cell death (20). Multiple pyroptotic pathways have been identified, each sensing different pathogen-associated molecular patterns (PAMPs) (24–27) or pathogen-associated aberrant activities (28, 29). The sensor converges on the activation of a cysteine-aspartic protease: caspase-1 for canonical inflammasome pathways (30, 31), and caspase-4/5 for later-discovered pathways designated noncanonical (32–34). Inflammatory caspases cleave and activate gasdermin D (35, 36), which assembles into pores in host plasma membranes (37). This leads to release of cytokines IL-18 and IL-1β (38), and total lysis for many cell types (39).

*P. aeruginosa* effectors ExoS and ExoT can reduce IL-1β secretion in macrophages (40–42), suggesting an ability to interfere with inflammasome activation. Responses of macrophages and epithelial cells can differ. While inflammasomes are well-studied in macrophages and other myeloid cells, the repertoire of inflammasomes in epithelial cells is usually limited, and their functions less-well studied (43). One inflammasome pathway studied in some corneal diseases is NLRP3 (44, 45), which can detect numerous stimuli (46) including ionic flux from pore formation (47), lysosomal damaging agents (48), and bacterial RNA (49). However, not all relevant studies verified which corneal cell type expressed NLRP3, or explicitly determined if NLRP3 was activated, versus other inflammasome pathways that could also yield mature IL-1β. Recently, both caspase-4, which detects cytoplasmic LPS (50), and NLRP1, which detects pathogenic enzymatic activities and dsRNA (28, 51), were shown to be expressed and functional in the human corneal epithelium (52, 53). The Protein Atlas RNAseq data set for the corneal epithelial cell line hTCEpi (54) provides evidence for expression of caspase-4, caspase-5 (an additional LPS sensor (33, 55)), NLRP1, and NLRC4 (although its required sensor protein NAIP (56) was not detected) (57). While NLRP3, Pyrin, and AIM2 inflammasomes were not detected in these cells, this does not preclude upregulation upon specific stimuli or in vivo (57).

Having shown that the T3SS effector ExoS inhibits rapid cell lysis induced by intracellular *P. aeruginosa* in corneal epithelial cells (19), we investigated the mechanisms underlying host cell death elicited and modulated by intracellular *P. aeruginosa*. In doing so, we also leveraged the finding that *P. aeruginosa* mutants lacking the entire T3SS (Δ*exsA* mutants) remain toxic to corneal epithelial cells (58), contrasting with HeLa and CHO cells (59, 60). Results showed that caspase-4 is required for rapid corneal epithelial cell death in response to *P. aeruginosa* invasion, and in this way limits accumulation of an intracellular population of bacteria. While caspase-4-dependent death occurred even if bacteria lacked the ability to leave vacuoles/enter the cytoplasm, the response was faster when they could. Moreover, the death response was enabled in otherwise unresponsive HeLa cells after stimulation with IFN-γ, which has been shown to limit cytoplasmic sub-populations of *Salmonella* in a manner dependent on caspase-4 and GBP proteins (61). Since ExoS inhibits the cell death response to intracellular *P. aeruginosa*, these results implicate ExoS targeting a caspase-4-dependent response as another contributor to its well-established role in intracellular survival by *P. aeruginosa*.

## Results

### *P. aeruginosa* lacking the T3SS kill corneal epithelial cells but not HeLa cells

Generally, it is thought that *P. aeruginosa* mutants missing the T3SS (i.e., Δ*exsA* mutants) do not kill epithelial cells, as shown for HeLa and CHO cells (59, 60). However, our published work has shown that Δ*exsA* mutants are able to kill corneal epithelial cells, with contributions made by the intracellular population (58). Recognizing that differences between cell types could assist in deciphering mechanisms, we compared corneal epithelial cells (hTCEpi) (54) to HeLa cells using the same methods. Impact of wild type PAO1, Δ*exsA* mutants (lacking the entire T3SS), and Δ*exoSTY* mutants (lacking all known T3SS effectors) were studied. After a 3-hour invasion period, extracellular bacteria were eliminated using the non-cell permeable antibiotic amikacin (3). Cell death rates were measured with a FIJI macro that counts propidium iodide-positive nuclei (permeabilized cells) and Hoechst-labeled nuclei (total cells) and reports a ratio over a 20-hour post-infection period (19). The results confirmed that the two cell types were differentially susceptible. Death rates for corneal epithelial cells were statistically similar between wild type and Δ*exsA* mutants (**Figure 1A-B**), whereas HeLa cells infected with Δ*exsA* mutants showed very little cell death even 20-h post infection (**Figure 1C-D**). In both cell types, the Δ*exoSTY* mutant yielded the most rapid cell death as noted previously (19). Thus, corneal epithelial cells have an intrinsic response to T3SS-null bacteria that is absent in HeLa cells, which may underly cell death occurring in response to T3SS-positive bacteria.

**Figure 1.**
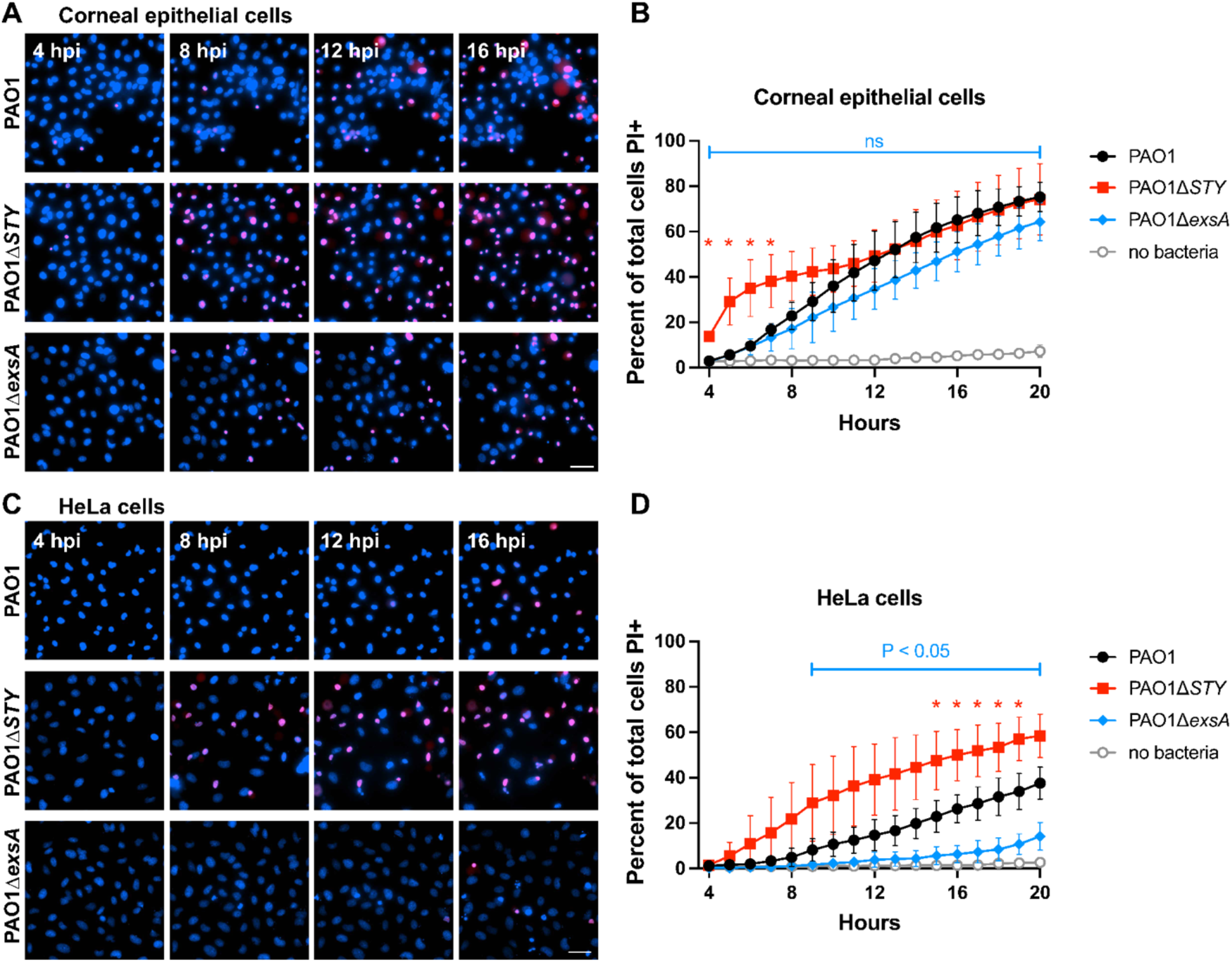
Host cell death rates from *P. aeruginosa* mutant infections. **(A)** Corneal epithelial cells (hTCEpi) were infected with indicated strain: wt PAO1, PAO1Δ*exoSTY*, or PAO1Δ*exsA* for 3 hours at an MOI equal to 10. Hoechst (blue) was added to label nuclei. Non-associated bacteria were removed, and media with amikacin and propidium iodide (red) was added to kill extracellular bacteria. Cells were imaged hourly from 4 to 20 hours post infection. Select fields from indicated times are shown. Scale bar = 50 µm. **(B)** The percent of cells positive for propidium iodide at each timepoint was determined by using a custom FIJI macro to segment all nuclei from the Hoechst channel, and all nuclei of dead cells in the propidium iodide channel. Multiple column t-tests were performed comparing wt PAO1 and PAO1Δ*exsA*, and SD error bars are displayed from three replicates. **(C)** Experiment was performed identically as in panel A, but using HeLa cells. Scale bar = 50 µm. **(D)** Analysis was performed as in panel B, using the data obtained in panel C.

### Dynamics of cell death in bacterially-occupied cells indicates roles for both T3SS machinery and T3SS effectors

Visual inspection of infected cells suggested that not all cells were occupied by bacteria, and that some non-occupied cells also died. Recognizing that this could skew results in bulk analyses of cell death rates, we performed additional experiments to analyze only bacterially-occupied cells. To enable this, we employed methods that allowed simultaneous detection of viable intracellular bacteria and host cell (19). Wild type (PAO1) was compared to mutants lacking the entire T3SS (Δ*exsA* mutants), and Δ*exoSTY* mutants lacking only T3SS effectors, using quantitative time lapse imaging over 20 h. Since Δ*exsA* mutants do not kill HeLa cells, this analysis was only done using corneal epithelial cells.

A method of detecting intracellular bacteria is needed, as amikacin-killed bacteria adhere to the dishes and cells, making intracellular bacteria impossible to distinguish if using constitutive fluorophore expression. To detect Δ*exsA* mutants inside cells, we previously used an arabinose-inducible green fluorescent protein (GFP) expression vector pBAD-GFP (58). This method involves induction of GFP expression using arabinose induction only after bacteria have invaded cells and extracellular populations are killed (using amikacin). To keep experimental methods consistent across samples, we explored the feasibility of using the induction method to also study wild type and the Δ*exoSTY* mutants. In prior studies, these strains were visualized using a reporter for the T3SS (3), because T3SS effectors are needed for intracellular replication (5, 19). Unfortunately, fewer bacterial cells exhibited cytoplasmic spread using the arabinose induction method compared to the T3SS reporter method (**Supplemental Figure 1A**). Moreover, while the host cell death rate was similar for wild type PAO1 using both methods, it trended lower for Δ*exoSTY* mutants expressing arabinose-induced GFP (**Supplemental Figure 1B)**. This suggested GFP induction impacts the T3SS or some other factor relevant to cytoplasmic entry or spread: a critical step for intracellular infection by wild type and Δ*exoSTY* mutants, but not for Δ*exsA* mutants (which remain in vacuoles (58)). Indeed, T3SS effector secretion was reduced *in vitro* using EGTA stimulation in the presence of arabinose when bacteria were also transformed with pBAD-GFP (**Supplemental Figure 1C**). Thus, the T3SS expression plasmid pJNE05 was used to study intracellular wild type and Δ*exoSTY* mutants, reserving arabinose induction of GFP for the Δ*exsA* mutant. A limitation of this approach is that a sub-population of intracellular wild type bacteria can remain T3SS-negative (58) and would be undetected in this analysis. However, the advantage of this approach is that T3SS-dependent outcomes for intracellular bacteria can be spatially isolated and examined independently, and then compared to homogenous populations of Δ*exsA* mutants missing the T3SS apparatus.

Representative images from a time-lapse experiment using Δ*exsA* mutants are shown in **Figure 2A**. As expected, Δ*exsA* mutants localized to vacuoles inside cells (58). Most bacteria-occupied cells died before the end of the 20 h assay, as shown by propidium iodide labeling of the nucleus. However, **Figure 2B** shows an invaded cell still alive at 20 hours (no propidium iodide labeling), representing a minority of cells at this time point. Videos of these time lapse experiments are available in **Supplemental Movie 1**, which includes cells visualized in **Figures 2B and 2C** at 20 hours post infection. As shown previously, both wild type PAO1 and Δ*exoSTY* mutant infected cells (shown in **Figure 2D)** replicated in the host cell cytoplasm after bacterial escape from vacuoles (19). In both cases, most invaded cells died by the end of the assay, the Δ*exoSTY* mutant doing so even more rapidly, expected due to lack of ExoS which normally counters cell death (19). Hours following host cell death, propidium iodide of intracellular bacterial bodies eventually replaced GFP signal (**Figure 2C and Supplemental Movie 1)**.

**Figure 2.**
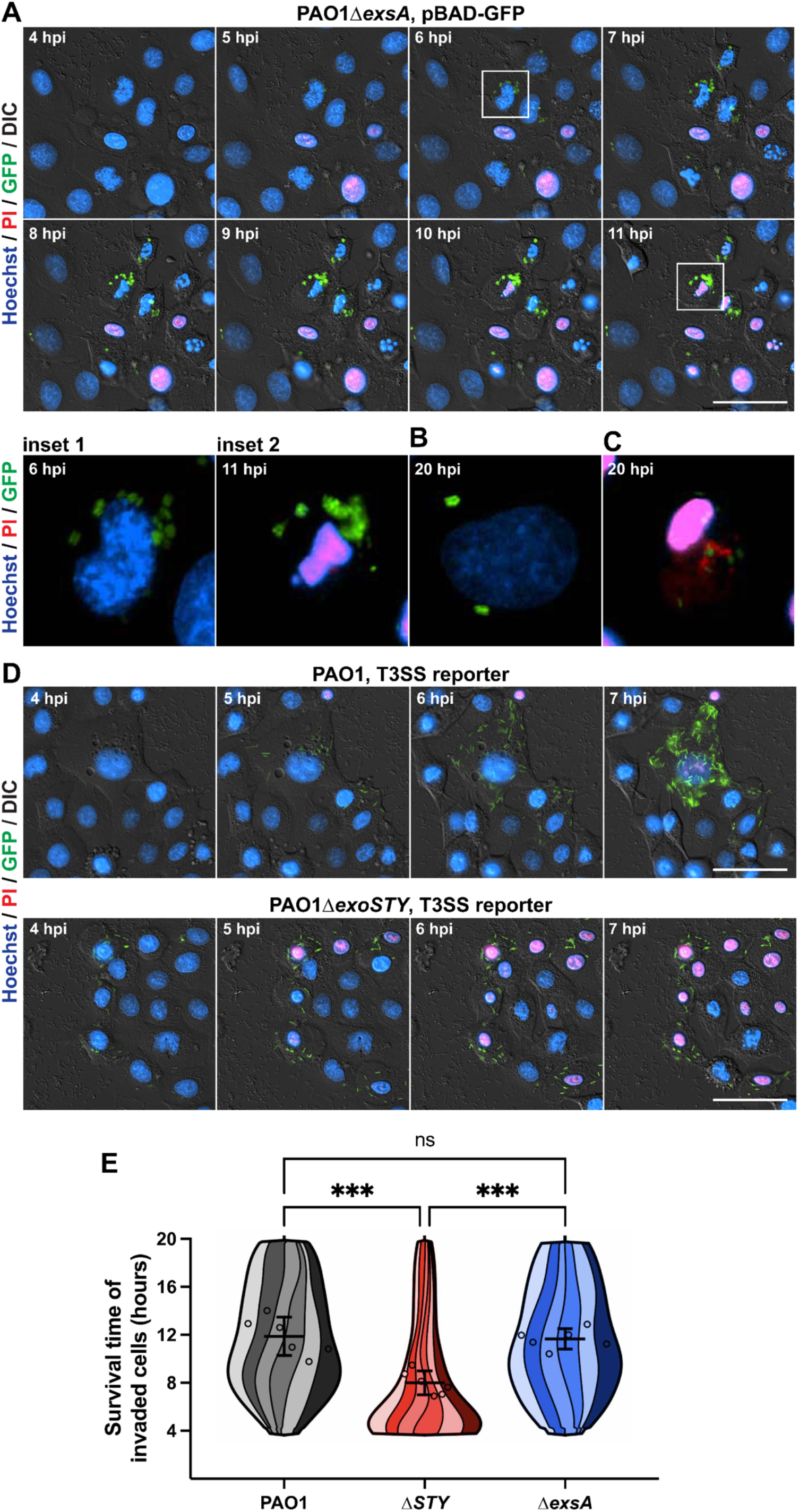
Intracellular Δ*exsA* mutants kill invaded cells at a similar rate to wt PAO1-invaded cells. **(A)** Bacteria were transformed with an arabinose-inducible GFP plasmid. Corneal epithelial cells (hTCEpi) were infected at an MOI of 10 and extracellular bacteria eliminated at 3 hours post infection using amikacin. A final concentration of 1% arabinose was used to induce GFP in surviving intracellular bacteria beginning at 3.5 hours post infection. Time lapse imaging was conducted hourly from 4 to 20 hours, and host cell nuclei detected with Hoechst and propidium iodide to determine time of death. Scale bar = 50 µm. Insets 1-2 show the boxed cell at 6 or 11 hours post infection, respectively. **(B)** An example of a live invaded cell from 20 hours post infection. **(C)** Example of a dead invaded cell from 20 hours post infection, where the PI channel has been saturated such that PI-positive bacteria can be visualized. Scale bar = 50 µm. **(D)** The T3SS-GFP reporter pJNE05 was used to visualize wt PAO1 and PAO1Δ*exoSTY* infections, as described previously (19), to compare to PAO1Δ*exsA* infections. Scale bar = 50 µm. Time lapses from panels A through D are available as **Supplemental Movie 2**. **(E)** A computational analysis approach was used to segregate only invaded cells, and determine their survival time in hours. PAO1 condition includes 881 cells over six replicates; PAO1Δ*exoSTY*, 967 cells; PAO1Δ*exsA*, 862 cells. The survival times of all invaded cells were combined a single super violin plot. Mean survival times with SD error bars are displayed, and significance determined by One-way ANOVA. *** P < 0.005.

A computational approach was used to measure when populations of invaded cells died (19), and a super violin plot visualization script developed by Kenny and Schoen was used to show independent replicates as stripes single plots (62). This analysis examined 2,710 total invaded cells over 6 replicate experiments. Results confirmed that intracellular Δ*exsA* mutants killed their occupied cells at a broad range of time points, and the mean value of invaded cell survival times was unexpectedly similar to wild type PAO1 (**Figure 3E**). The analysis also confirmed that the intracellular Δ*exoSTY* mutant infections caused cell lysis at a more rapid rate, significantly different from both wild type PAO1 and Δ*exsA* mutants (19). Thus, while cell death can be driven by vacuolar contained bacteria, it was more rapid if bacteria could access the cytoplasm while lacking effectors.

**Figure 3.**
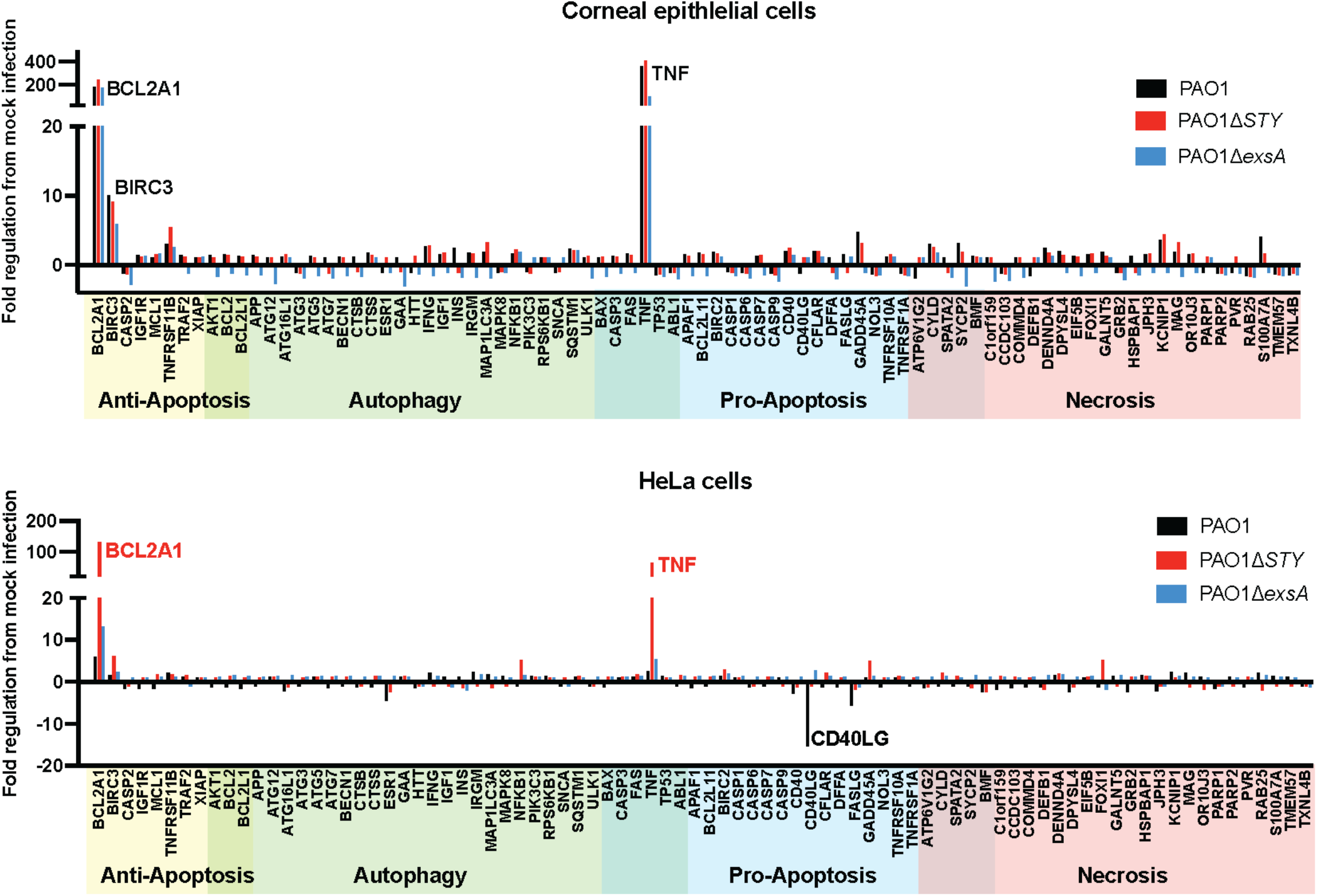
Transcriptional responses of infected host cells. **(A)** Corneal epithelial cells (hTCEpi) were infected with wt PAO1, PAO1Δ*exoSTY*, or PAO1Δ*exsA* for 3 hours at an MOI equal to 10. Non-associated bacteria were removed, and media with amikacin was added for 1 additional hour. At four hours, RNA was purified from infected cells, and RT PCR analysis performed using a commercial array for genes associated with specific cell death pathways. **(B)** Experiment performed identically to panel A, except using HeLa cells.

### T3SS-null mutant bacteria engage NF-κB signaling in corneal epithelial cells.

To explore molecular mechanisms involved in *P. aeruginosa*-driven cell death, a real time PCR array with genes involved in regulated cell death pathways was used to study candidate host responses. Results showed that TNFα and BCL2A1 were upregulated in excess of fifty-fold in both cell lines: (**Figure 3**). In the corneal cells, wild type and both mutants upregulated these genes, while in HeLa cells, only the PAO1Δ*exoSTY* mutant had that impact.

Since TNFα and BCL2A1 can be regulated by NF-κB, activation of NF-κB was measured by observing translocation of p65 to the nucleus (**Figure 4A**). In corneal epithelial cells, infection with each tested strain caused p65 translocation, correlating with positive target gene transcription. In HeLa cells, only infection with Δ*exoSTY* mutants led to p65 translocation (**Figure 4B)**. The result with wild type PAO1 infection was difficult to discern due to cell rounding and low cytoplasmic area; however, p65 appeared excluded from the nuclear region of HeLa cells (**Figure 4B, inset)**. Quantification of p65-positive nuclei was accomplished with an ImageJ macro (see methods) (**Figure 4C and D**). This finding correlated with the transcriptional data, and is suggestive of a mechanism in which the T3SS effectors limit p65 translocation in HeLa cells, but not corneal epithelial cells. Since TNFα is associated with both apoptotic and necrotic cell death, TNFα secretion from infected cells was measured by ELISA. The results showed that while wild type PAO1 stimulated TNFα secretion from corneal epithelial cells, higher levels were detected after infection with the Δ*exoSTY* mutant, and the Δ*exsA* mutant causing an intermediate response (**Figure 4E**). TNFα was unable to be detected from HeLa cells infected with any strain.

**Figure 4.**
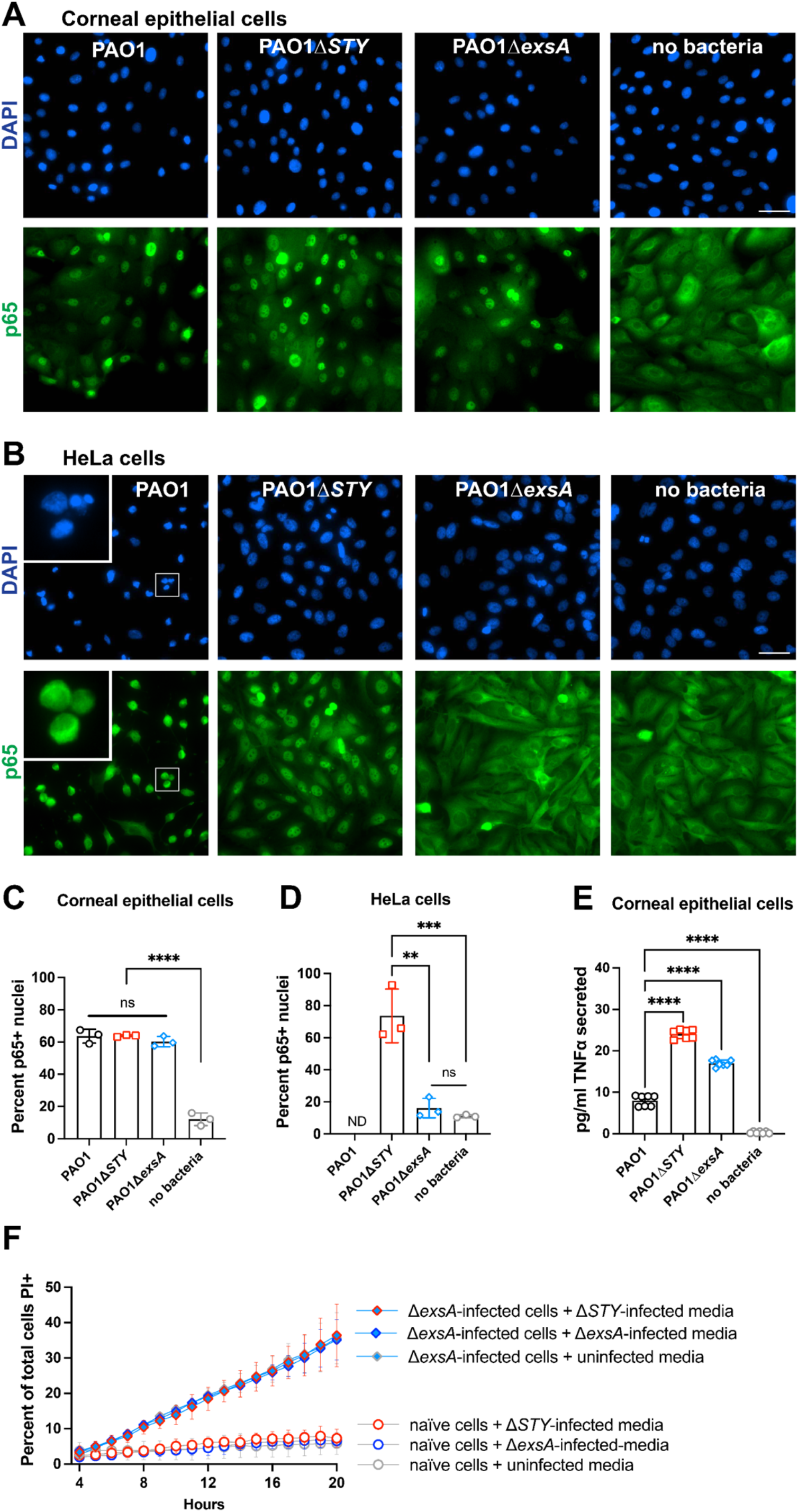
NF-κB signaling and the role of secreted factors in cell death. **(A)** Corneal epithelial cells (hTCEpi) were infected with wt PAO1, PAO1Δ*exoSTY*, or PAO1Δ*exsA* for 3 hours at an MOI equal to 10. Then media was replaced with amikacin-containing media. Cells were fixed at 4 hours post infection and p65 localization was determined using immunofluorescent staining. Nuclei were labeled with DAPI. Scale bar = 50 µm. **(B)** Experiment performed identically to panel A, using HeLa cells. **(C)** Quantification of p65-positive nuclei from corneal epithelial cells, and (**D**) HeLa cells. PAO1-infected HeLa cells were not able to be accurately quantified (indicated using “ND”) by automated analysis due to the small and irregular cytoplasmic area. (**E**) Corneal epithelial cells were infected as described in panel A. Supernatant of infected cells was collected at 4 hours post infection and TNFα measured by ELISA. **(D)** Supernatants of corneal epithelial cells infected with PAO1Δ*exoSTY*, PAO1Δ*exsA*, or uninfected were collected at 4 hours post infection and sterilized with 0.22 µm filter, and used as the replacement media on uninfected cells or PAO1Δ*exsA* infected cells at 3 hours post infection, combined with amikacin and propidium iodide. Hoechst was used to label all nuclei. The cell death rates were measured by time lapse imaging, error bars display SD from three replicates.

Having found TNFα secretion levels correlated with cell death rates, we next asked if there was a direct causative relationship that might also explain why some cells without visible intracellular bacteria could also be killed. Thus, media from naïve cells or Δ*exsA* mutant-infected cells was replaced with filter-sterilized media from Δ*exoSTY* mutant-infected cells (high TNFα). The outcome showed that this did not enhance host cell death rates (**Figure 4F**), suggesting cell death is not driven by a secreted factor such as TNFα, and requires bacteria present.

### Caspase-4 is involved in cell death response to intracellular *P. aeruginosa*

The morphology of invaded cell death included membrane integrity loss, intact nucleus, and transient blebbing, all suggestive of pyroptosis (**Figure 2**) (22). Since caspase-4 can restrict numerous other Gram-negative bacteria (61, 63), and the pathogen *Shigella* inhibits the caspase-4 inflammasome by multiple mechanisms to survive (64–66), and corneal epithelial cells were shown to express caspase-4 (53), we tested the hypothesis that caspase-4 was involved in driving cell death in response to intracellular *P. aeruginosa*. We generated *CASP4* knockout corneal epithelial cells, and loss of caspase-4 protein was confirmed by Western blot (**Supplemental Figure 2**). Control cell lines with non-targeting guide RNA were also generated using an identical selection strategy and grown up as monoclonal lines; each exhibited death rates consistent with wild type, non-transduced hTCEpi cells (**Supplemental Figure 3**).

Invasion by wild type PAO1 yielded similar cell death timing in both *CASP4* knockout cells as wild type corneal epithelial cells (**Figure 5A, B, C**). In contrast, Δ*exoSTY* mutant-invaded cells experienced significantly increased survival times (**Figure 5A, B, D**), which allowed substantial accumulation of intracellular bacteria. A smaller but significant increase of invaded cell survival times were also observed with PAO1Δ*exsA* mutant-occupied *CASP4* knockout cells (**Figure 5E**). Bacterial replication was measured using summed GFP area contained in live cell masked region generated by the FIJI macro (19). Results indicate increased bacterial replication in *CASP4* knockout cells, with the most substantial increase observed in cells invaded by Δ*exoSTY* mutants (**Figure 5F-H**). These data implicated the non-canonical inflammasome as a dominant response to cytoplasmic invasion of T3SS-positive *P. aeruginosa* independently of the T3SS effectors, and also show involvement in responding to vacuole-occupying bacteria. Unchanged cell death timing for wild-type PAO1 in *CASP4* knockout cells aligns with our previously published data showing that the T3SS effector exotoxins, specifically the ADP ribosyltransferase activity of ExoS (19), counters cell death. Of note, while *CASP4* knockout reduced corneal epithelial cell death associated with intracellular Δ*exsA* mutants, many of the cells still died by the end of the 20-hour imaging time frame (**Figure 1 and 5E**). This implies additional mechanisms for recognizing vacuolar/T3SS-negative *P. aeruginosa* in addition to caspase-4 which are also absent from HeLa cells.

**Figure 5.**
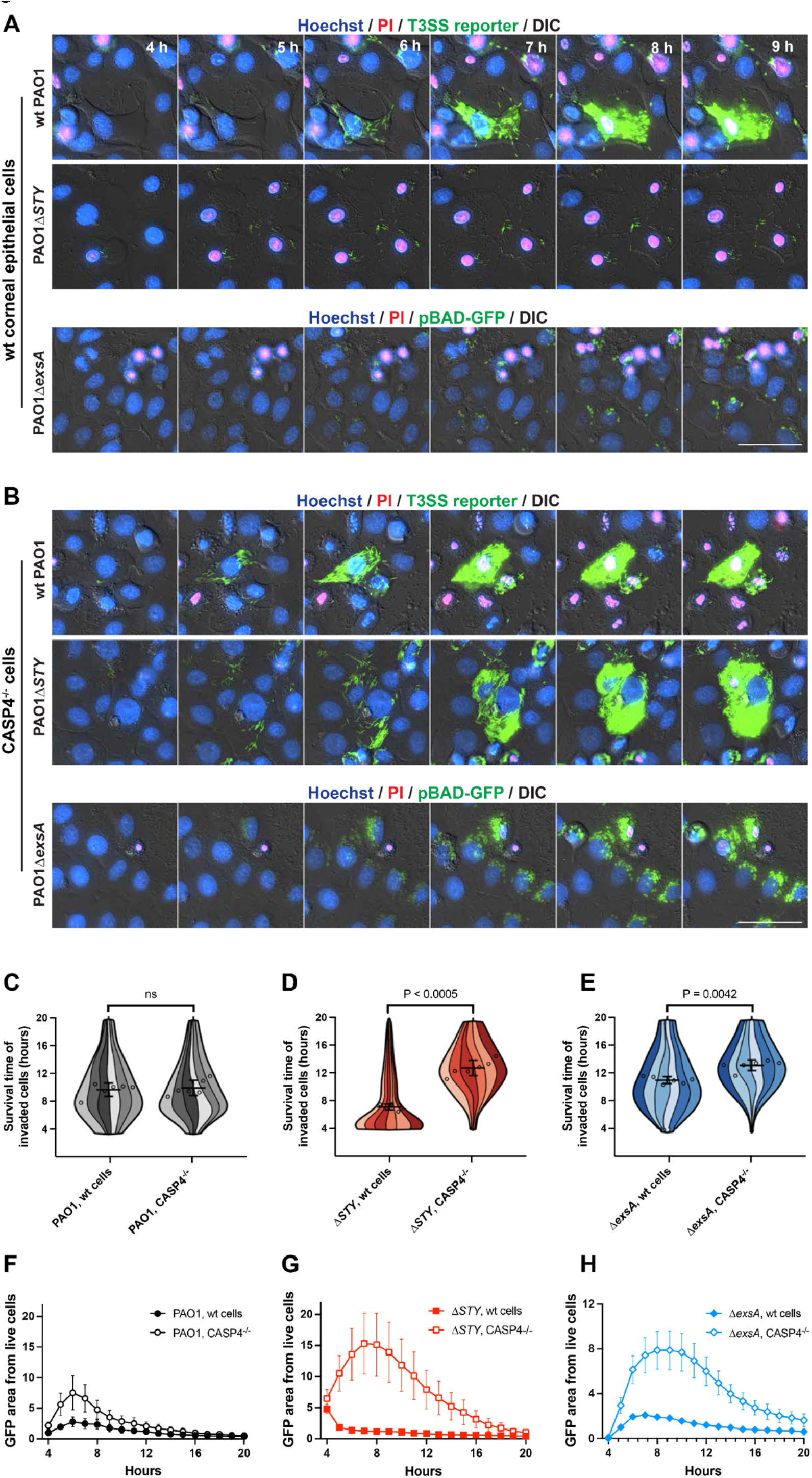
Caspase-4 limits intracellular replication and persistence of T3SS mutant bacteria. (A) Corneal epithelial cells (hTCEpi) or (B) cells knocked out for caspase-4 were infected with wt PAO1, PAO1Δ*exoSTY* (each visualized by T3SS-GFP reporter) or PAO1Δ*exsA* (pBAD-GFP induced) at an MOI of 10. Nuclei were labeled with Hoechst and propidium iodide. Extracellular bacteria were eliminated with amikacin at 3 hours post infection, and imaged hourly from 4 to 20 hours post infection. Images from 4-9 hours post infection are shown. Scale bar = 50 µm. Full-field time lapses are available as **Supplemental Movie 3**. (**B-D**) (C-D) Cells from indicated infections were analyzed with a computational approach to segregate only invaded cells and measure survival times in hours. Six replicates were combined into a single super violin plot. Mean survival times with SD error bars are displayed, and significance determined by Student’s t-test. (F-H) GFP area was summed within the boundaries of live cells using the same data set as panels C-E, and normalized to PAO1 (wt) at 4 hours post infection.

Prior studies implicate Exotoxin S in delaying host cell lysis in response to invasion (19), thus we examined the Δ*exoS* mutant in both wt corneal epithelial cells and *CASP4* knockout cells to evaluate whether there is a role for ExoT and ExoY in suppressing caspase-4-mediated lysis or supporting cytoplasmic bacterial grown (**Supplemental Figure 4**). Comparing to the aforementioned study which tested strains encoding a single effector (e.g., Δ*exoST* or Δ*exoSY*) (19), the ΔexoS mutant showed a similar phenotype to Δ*exoSY* mutant, which expresses only ExoT: bacteria were unable to replicate substantially within in wt corneal epithelial cells (**Supplemental Figure 4**). Bacterial mutant replication was rescued in *CASP4* knockout cells. However, the timing of Δ*exoS*-invaded wt cell survival was irregular across the timeframe observed (**Supplemental Figure 4C**), which may relate to ExoT’s ability to block invasion (67, 68), or interfere with other inflammasomes (41). However, ExoT and ExoY were unable to contribute to intracellular replication in cells with caspase-4.

### IFN-γ-stimulation enables HeLa cells to respond to intracellular *P. aeruginosa*

HeLa cells support cytoplasmic hyper-replication of *P. aeruginosa* before significant impact on host cell viability occurs (3, 19). The bacterium *Salmonella* accomplishes a similar feat (69). Santos and colleagues demonstrated that IFN-γ stimulation causes HeLa cells to lyse in response to *Salmonella* entry into the cytosol, which was associated with upregulation of GBP proteins and stimulation of the caspase-4 inflammasome. (61). We examined the impact of IFN-γ stimulation on HeLa cells containing intracellular *P. aeruginosa* (**Figure 6**). The kinetics of wild type PAO1-induced host cell death remained unchanged. However, the rate of cell death increased for both PAO1Δ*exoSTY* and for PAO1Δ*exsA* infections (**Figure 1 and 6**) (59). IFN-γ-treated HeLa cells occupied by Δ*exsA* mutants died over varied timepoints during the 20 hours of infection similar to results for corneal epithelial cells. Aligning with the quick timing of cell lysis, PAO1Δ*exoSTY* mutants were no longer able to replicate effectively in the cytoplasm of IFN-γ-treated HeLa cells. Thus, IFN-γ-treated HeLa cells resembled corneal epithelial cells in their response to intracellular *P. aeruginosa*.

**Figure 6.**
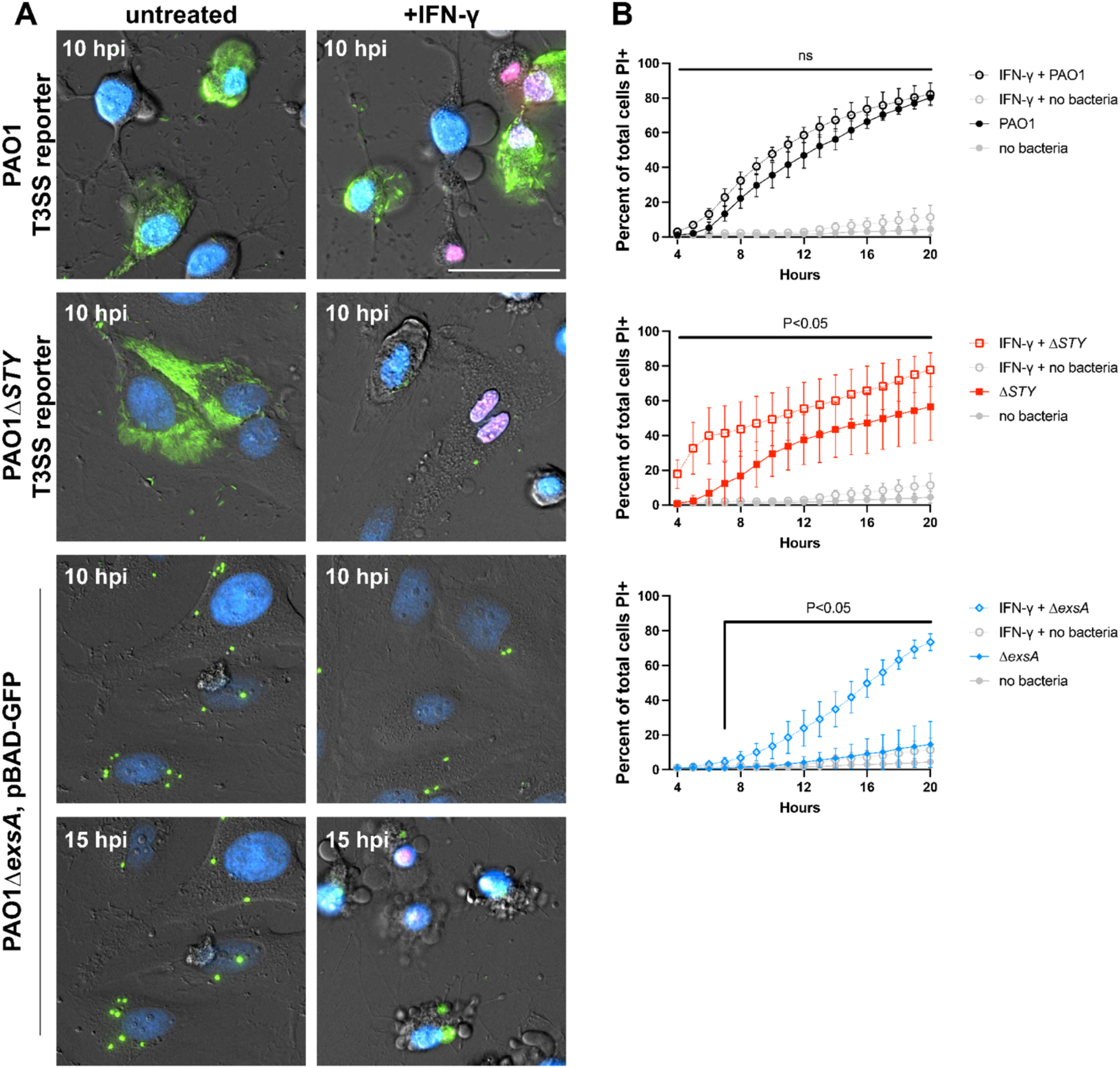
IFN-γ-stimulated HeLa cells are not permissive for PAO1Δ*exoSTY* intracellular replication nor PAO1Δ*exsA* intracellular persistence. (A) HeLa cells were treated with 50 ng/ml IFN-γ for 16 hours prior to infection with wt PAO1, PAO1Δ*exoSTY* (each using the T3SS-GFP reporter) or PAO1Δ*exsA* (pBAD-GFP induced) at an MOI of 10. Nuclei were labeled with Hoechst and propidium iodide. Extracellular bacteria were eliminated with amikacin at 3 hours post infection, and imaged hourly from 4 to 20 hours post infection. Select time frames are shown at either 10 or 15 hours post infection. Scale bar = 50 µm. Full-field time lapses are available as **Supplemental Movie 4**. **(B)** The rate of cell death was determined using host cell nuclear stains. All panels are from the same experiment with only indicated bacterial strain shown. Error bars show SD from three replicates. Multiple column t-tests were performed comparing IFN-γ-treated cells to untreated.

## Discussion

Invasive (ExoS-expressing) strains of *P. aeruginosa* can thrive inside a wide range of host cells (2, 3, 5). After invading an epithelial cell, ExoS plays multiple roles in supporting intracellular survival and replication, including inhibition of host cell lysis (19). This study aimed to understand the underlying host response inhibited by ExoS. Knowing *P. aeruginosa* diversifies intracellularly into vacuolar and cytoplasmic populations, bacteria in both locations trigger cell death, their locations are determined by T3SS expression state, and that host cell lysis can be suppressed by ExoS, we used T3SS mutants to interrogate mechanisms. Results showed that cell death in response to cytoplasmic *P. aeruginosa* lacking T3SS effectors involved caspase-4; its deletion inhibiting rapid pyroptotic lysis. Deletion of caspase-4 also partially alleviated cell death caused by bacteria lacking the T3SS that were restricted to vacuoles (Δ*exsA* mutants). We further found that while HeLa cells lack this response to intracellular *P. aeruginosa*, it could be induced by IFN-γ, a stimulator of numerous genes including factors that assist in activation of the caspase-4 inflammasome (61). This suggests one difference between corneal epithelial cells and HeLa cells may be among interferon stimulated genes (such as GBP1-4), or the ability of corneal epithelial cells to promptly upregulate inflammatory genes in response to *P. aeruginosa* (**Figure 3**). Since ExoS inhibits cell death in response to cytoplasmic bacteria (19), these findings implicate ExoS-mediated interference with the non-canonical inflammasome pathway. ExoS is only capable of prolonging cell survival to the same level as Δ*exsA* mutant-invaded cell death, suggesting its role is restricted to the host response to cytoplasmic bacteria.

Since *P. aeruginosa* is often assumed an extracellular pathogen, previous studies of host epithelial cell death have been from the perspective of extracellular bacteria. In that regard, ExoS alone has cytotoxic activities. HeLa cells can undergo apoptotic cell death based on both caspase-3 (59) and caspase-8 activity (70), and activation of apoptosis through JNK1 and cytochrome c release (71). This was confirmed using a mammalian expression system for introducing ExoS in the absence of bacteria, which showed it was sufficient to induce HeLa cell apoptosis even in the absence of bacterial interaction (72). Placing our findings into the context of these earlier HeLa cell studies, we note differences in our experimental systems: while they explored mechanisms for cell death caused by ExoS, we studied cell death triggered by live intracellular bacteria lacking ExoS. Were intracellular bacteria present in these assays, HeLa cells would not have responded to them without stimulation. Our experiments also provide an analysis of only invaded cells excluding extracellular bacteria to focus on impact of intracellular bacteria, and used corneal cells which naturally respond to bacterial pathogen-associated molecular patterns. We then showed that IFN-γ-stimulated HeLa cells respond in the same manner: ExoS inhibits rather than drives a host cell death response. In sum, while introducing ExoS in isolation is a useful surrogate for *in vitro* experimentation to address specific questions, other factors modify the impact of ExoS in other situations. Despite differences in experimental models, it remains important to reconcile outcomes. The innate ability of ExoS to both drive and inhibit cell death, involving different types of regulated cell death, and correlating with presence or absence of internalized *P. aeruginosa,* is interesting and worth exploring. Indeed, it could aid in determining mechanisms by which ExoS blocks pyroptosis, given crosstalk between pyroptosis and apoptosis (73).

The specific mechanism by which the ADPr activity of ExoS interferes with caspase-4-mediated cell lysis remains to be identified. Recently, *Shigella flexneri* was shown to block the non-canonical inflammasome by multiple mechanisms: caspase-4 inactivation from ADP-riboxanation by the effector OspC3 (64, 65), ubiquitylating gasdermin D (74), and GBP1 (75) to target each for degradation. ExoS is unique among bacterial ADP ribosyltransferases for having numerous host substrates, though currently none are known to interact with the caspase-4/11 pathway (76). Since there are many possibilities to investigate, comprehensive identification of uncharacterized ExoS substrates and their potential involvement will require a separate study. Considering that *CASP4* knockout does not fully prevent cell death triggered by vacuole-bound T3SS mutants (**Figure 6**), it would also be worth exploring which additional cell death pathways are involved.

In summary, the results of this study demonstrate a role for caspase-4 in limiting intracellular colonization of *P. aeruginosa* in ocular surface cells, a phenomenon triggered by the presence of intracellular bacteria but inhibited by the T3SS effector ExoS. Importantly, the data support the notion that *P. aeruginosa* can function as both an intracellular and extracellular pathogen, suggestive of a T3SS effector function devoted to inhibition of an *intracellular* pattern recognition receptor, consistent with its unusual capacity to adapt to its environment, survive hardship, and exist ubiquitously. Ultimately, development of effective strategies to prevent or treat the devastating infections caused by *P. aeruginosa* will require a thorough appreciation of these complexities.

## Methods

### Bacterial strains and cell lines

*P. aeruginosa* strain PAO1 and isogenic mutants of *exsA*, or ExoS/ExoT/ExoY, or ExoS were used for all infection experiments (77). Plasmids used for visualization were pJNE05 (78) and pBAD-GFP (58). Plasmid selection was achieved with 100 µg/ml gentamicin. Corneal epithelial cells (hTCEpi) (54) were maintained in KGM-2 media (Lonza) lacking gentamicin. HeLa cells were maintained in phenol red-free DMEM (Gibco) with 10% FBS.

### CRISPR-Cas9 knockout cell line

Lentiviral particles for transduction were generated from 293T cells transfected with psPAX2 (Didier Trono, Addgene plasmid # 12260) and pMD.2 (Didier Trono, Addgene plasmid # 12259), combined with either Cas9 expression vector plenti-Cas9-blast (Feng Zhang, Addgene plasmid # 52962) (79) or a guide RNA vector plenti-LIC-Puro which was adapted for ligation-independent cloning (gifted by Moritz Gaidt) (80). Guide RNA sequences targeting caspase-4 (5’-CCACAGAAAAAAGCCACTTA - 3’) or a non-targeting control sequence (5’-GACGGAGGCTAAGCGTCGCA - 3’) were cloned into plenti-LIC-Puro using ligation independent cloning. After six hours, media on 293T cells was replaced with KGM-2. Media was collected at 48 hours and passed through a 0.45 µm filter and added directly to hTCEpi cells, which were then centrifuged at 1200g for 90 minutes at 37°C. Antibiotic selection with blasticidin (50 µg/ml) or puromycin (10 µg/ml) was performed after two days. For generation of monoclonal lines, single cells were seeded in 96-well plates. Isolated colonies were identified and grown up over approximately 12 days before transfer into 6-well plates to avoid differentiation induced by crowding. *CASP4* knockout candidates were screened by Western blot: lysates from a 6-well plate were collected in 150 µl RIPA buffer, frozen and thawed, and then clarified by centrifugation for 20 minutes at 4°C. Samples were run on 10% stain free gels (Bio-Rad) and total protein visualized as a loading control prior to transfer to 0.2 µm PVDF membrane. Membrane was blocked with Bio-Rad EveryBlot block and probed with anti-caspase-4 antibody (Santa Cruz, sc-56056, 1:1000). Blot was washed using TBST, and antibody detected with goat-anti-rabbit HRP (Bio-Rad, 1706515, 1:5000).

### Infection experiments

One day preceding experiments, hTCEpi cells were plated on No. 1.5 glass-bottom 24 well plates (MatTek) in KGM-2 at 75% confluence with 1.15mM calcium to induce differentiation (54). HeLa cells were maintained in DMEM with 10% FBS and plated at 50% confluence on optical plastic 8-well chambered coverslips (ibidi). Where indicated, 50 ng/ml IFN-γ (Peprotech) was added at 16 hours prior to infections of HeLa cells and maintained throughout the infection and imaging.

Bacterial suspensions were made in PBS from 16-hour lawns grown on TSA media at 37°C. The absorbance at 540 nm was measured, and MOI of 10 was calculated using OD540 of 1 equal to 4 x 10^8^ CFU per mL. Bacteria were directly to cell culture media and allowed to infect for 3 hours. For live imaging, Hoechst (0.8 µg/ml) was added to the cells prior to infections and allowed to label the cells during the 3-hour infection period. Media was replaced with media containing the antibiotic amikacin (200 µg/ml) and propidium iodide (0.8 µg/ml). Where needed, a 1:10 dilution of 10% arabinose in media was added at 3.75 hours post infection.

### RT-PCR arrays

Cells in 10cm dishes were infected with PAO1, PAO1Δ*exoSTY*, or PAO1Δ*exsA* for 3 hours at an MOI of 10, and media was replaced for 1 hour with 200 µg/ml amikacin. At 4 hours, cells were washed twice in PBS and collected in 1ml TriReagent (Sigma). RNA was purified using Zymo Direct-zol RNA MiniPrep. cDNA synthesis was performed with the RT2 first strand kit (Qiagen) using 0.5µg RNA. RT PCR was performed on a Roche LightCycler 96 using the The RT² Profiler™ PCR Array Human Cell Death PathwayFinder (PAHS-212Z, Qiagen)w, and analyzed in Qiagen’s Data Center Analysis Portal. (https://geneglobe.qiagen.com/us/analyze).

### Fixed immunostaining

Cells were seeded on No. 1.5 glass coverslips and infected as described above. At four hours post infection, cells were washed twice in PBS and fixed in 4% paraformaldehyde in PBS for 10 m. Cells were washed twice in PBS, and neutralized with 150 mM glycine in PBS for 10 m. Cells were washed twice in PBS, and permeabilized/blocked (5% FBS, 2.5% cold fish skin gelatin, 0.1% TritonX-100, 0.05% Tween-20 in PBS) for 1 h. Antibody in solution (2.5% FBS, 1.25% cold fish skin gelatin, 0.1% TritonX-100, 0.05% Tween-20 in PBS) was incubated overnight at 4°C. Cells were washed 4 times (5 minutes), and secondary antibodies were added in antibody solution for 1 h. Cells were washed once, labeled with DAPI for 5 minutes, and washed twice (5 minutes). Coverslips were mounted using ProLong Diamond.

### Microscopy

Images for Figs 1 and 4A, B were captured on a Nikon Ti-E inverted microscope equipped with a Lumencor Spectra X LED Light Engine illumination source. All other images were captured on a Ti2-E inverted microscope with X-Cite XYLIS XT720S Broad Spectrum LED Illumination System. Both systems used Nikon perfect focus, an Okolab stage-top incubation chamber, a DS-Qi2 CMOS camera, and CFI Plan Apochromat Lambda D 40X air NA 0.95 objective. Time lapse fields were selected between hours 3 and 4 without observing fluorescence channels. Eight fields per condition were imaged hourly from 4 to 20 hours post infection.

### Image Analysis

Time-lapse images were computationally analyzed using two custom macros written for the FIJI package of Image J aspreviously described (19). The code for measuring rates of cell death for the whole population, and set of macros for tracking the invasion state of cells is available in a GitHub repository: https://github.com/Llamero/Nuclei_analysis-macro. Subsequent Python scripts for exporting the analysis TIF file and processing are available in the following GitHub repository: https://github.com/abbykroken/cell_survival_with_bacteria. An Image J macro for measuring a ratio of p65 stain intensity within the nucleus and the periplasmic region has been made available in the following GitHub repository: https://github.com/abbykroken/Nucleus-to-cell-ratio.

### In vitro T3SS effector secretion

Bacteria were grown for 5 hours in 5ml TSB supplemented with 100mM MSG and 1% glycerol at 37°C with shaking at 200 rpm. EGTA (2mM) was added to induce T3 secretion. OD 540 nm readings were taken prior to supernatant protein concentration and used to normalize volumes. Bacteria were centrifuged at 12000g to clarify 1ml of supernatant. Proteins were concentrated using TCA precipitation: 250 µl of 100% cold TCA was added to 1 ml supernatant for 30 minutes and centrifuged at 14000g for 5 minutes. The pellet was washed with 1 ml cold acetone twice and suspended in 4x laemmli buffer normalized to starting OD (max volume 50 µl). Proteins were visualized using 10% stain-free gels (Bio-Rad).

### Statistics

Statistical analyses were performed and data presented using Graph Pad Prism 9. Super violin plots were prepared using scripts published by Kenny and Schoen (62) and output images aligned on axes generated in Graph Pad Prism 9. Data were shown as a mean ± standard deviation (SD) of 3-6 independent experiments unless otherwise indicated. Comparison of two groups was performed by Student’s t-test, three or more groups by One-way ANOVA with Tukey’s post-hoc analysis. Comparison between two groups for total cell death rates overtime was performed by multiple column t-tests for each timepoint, and the two samples that were compared are specified in the figure legends. In each instance, * P < 0.05, ** P < 0.01, *** P < 0.005 and **** P < 0.001.

## Acknowledgements

Thanks to Arne Rietsch (Case Western Reserve University, OH) for providing *P. aeruginosa* strains and mutants, Timothy L Yahr (University of Iowa, IA) for plasmid pJNE05, and Danielle M Robertson (University of Texas Southwestern, TX) for hTCEpi cells. Thanks to Benjamin E Smith (University of California, Berkeley, CA) for continued support of FIJI image analysis macros. We also thank Russell E Vance (University of California, Berkeley, CA) for helpful suggestions during the course of this project.

ARK, PSM, VN, DJE, and SMJF designed the experiments; ARK, KAK, PSM, and EJJ performed the experiments; ARK, KAK, PSM, VN, DJE, SMJF analyzed and interpreted the data; ARK, DJE, and SMJF wrote the manuscript; ARK, DJE, and SMJF supervised the study.

This work was supported by the National Institutes of Health R01EY011221 (SMJF), R01EY034239 (ARK), F32 EY025969 (ARK), DP2 AI154432 (PSM), P30 DK089507 (PSM), F32 EY029152 (VN), and the Mallinckrodt Foundation (PSM). The funding agencies had no role in the study design, data collection and interpretation, or decision to submit the work for publication.

---

**Supplemental Movie 1. Corneal epithelial cells invaded by *P. aeruginosa* and mutants.** Corneal epithelial cells (hTCEpi) were infected with wt PAO1, PAO1Δ*exoSTY* (each expressing GFP by T3SS reporter), or PAO1Δ*exsA* (inducible GFP) for 3 hours at an MOI equal to 10. Then media was replaced with amikacin-containing media. Arabinose was added to PAO1Δ*exsA* condition at 3.5 hours post infection. Imaging was conducted hourly from 4 to 20 hours.

**Supplemental Movie 2. *CASP4* knockout corneal epithelial cells invaded by *P. aeruginosa* and mutants.** Corneal epithelial cells (hTCEpi) or *CASP4* knockout cells were infected with wt PAO1, PAO1Δ*exoSTY* (each expressing GFP by T3SS reporter), or PAO1Δ*exsA* (inducible GFP) for 3 hours at an MOI equal to 10. Then media was replaced with amikacin-containing media. Arabinose was added to PAO1Δ*exsA* condition at 3.5 hours post infection. Imaging was conducted hourly from 4 to 20 hours.

**Supplemental Movie 3. IFN-γ-stimulated HeLa cells invaded by *P. aeruginosa* and mutants.** HeLa cells were treated with 50 ng/ml IFN-γ for 16 hours prior to infection with wt PAO1, PAO1Δ*exoSTY* (each using the T3SS-GFP reporter) or PAO1Δ*exsA* (pBAD-GFP induced) at an MOI of 10. Nuclei were labeled with Hoechst and Propidium iodide. Extracellular bacteria were eliminated with amikacin at 3 hours post infection, and imaged hourly from 4 to 20 hours post infection. Arabinose was added to PAO1Δ*exsA* condition at 3.5 hours post infection. Imaging was conducted hourly from 4 to 20 hours. IFN-γ was maintained during infection and imaging.

**Supplemental Figure 1.**
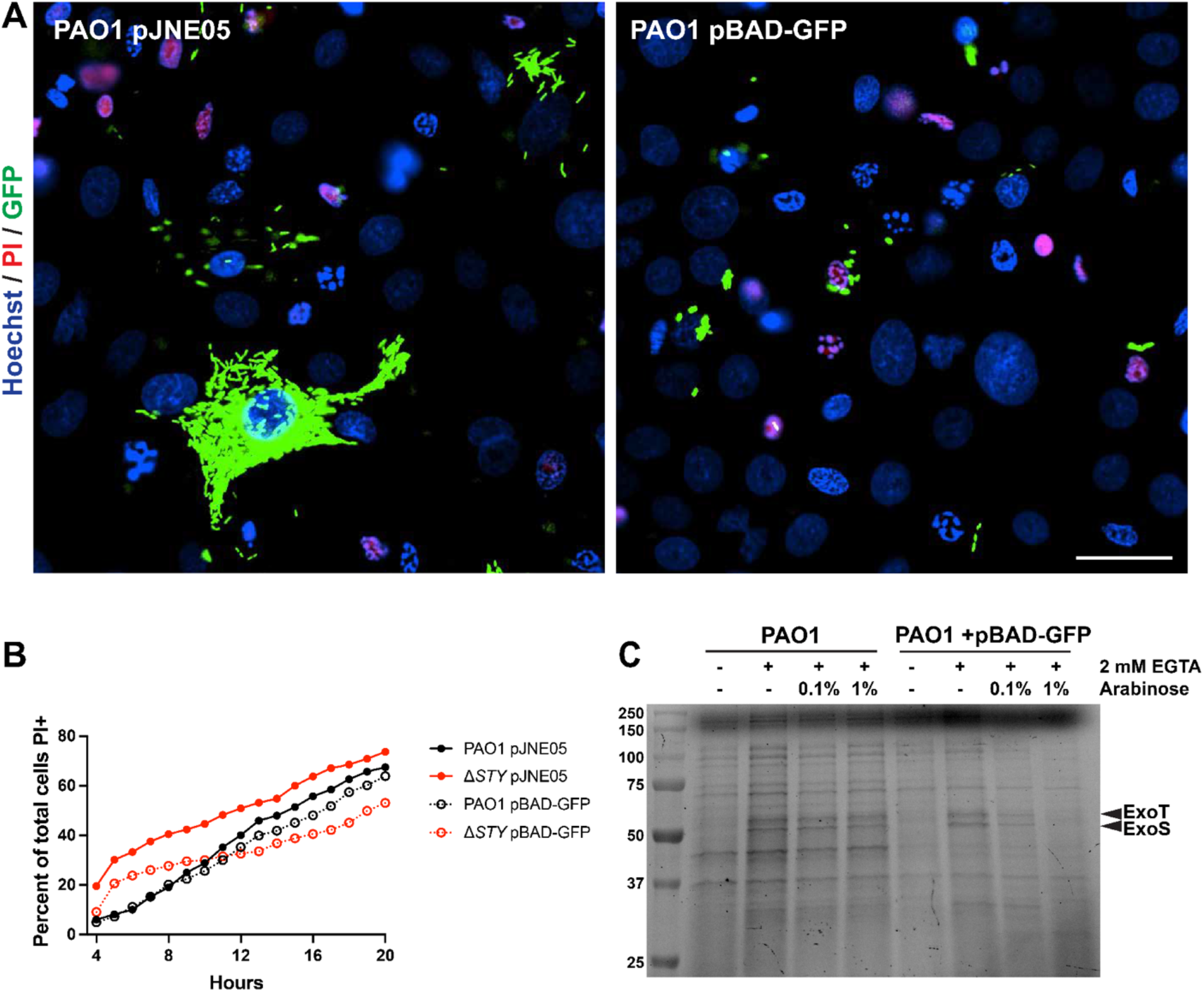
GFP expression from plasmid pBAD-GFP suppresses the T3SS. (A) wt PAO1 was transformed with the T3SS reporter plasmid pJNE05, or the GFP inducible plasmid pBAD-GFP and imaged hourly in time lapse. Example fields are shown in which bacteria expressing GFP under the T3SS reporter exhibit intracellular spread and replication, whereas those with inducible GFP show only limited intracellular replication. (B) The cell death rates of wt PAO1 or PAO1Δ*exoSTY* expressing GFP by either plasmid were measured, showing a trend of reduced cell killing rates for PAO1Δ*exoSTY* when GFP expression is induced. (C) wt PAO1 with indicated plasmid was grown for 5 hours in LB media with EGTA to induce the T3SS, and arabinose to induce GFP. Supernatant proteins were concentrated by TCA precipitation, and the bands constituting ExoS and ExoT are indicated.

**Supplemental Figure 2.**
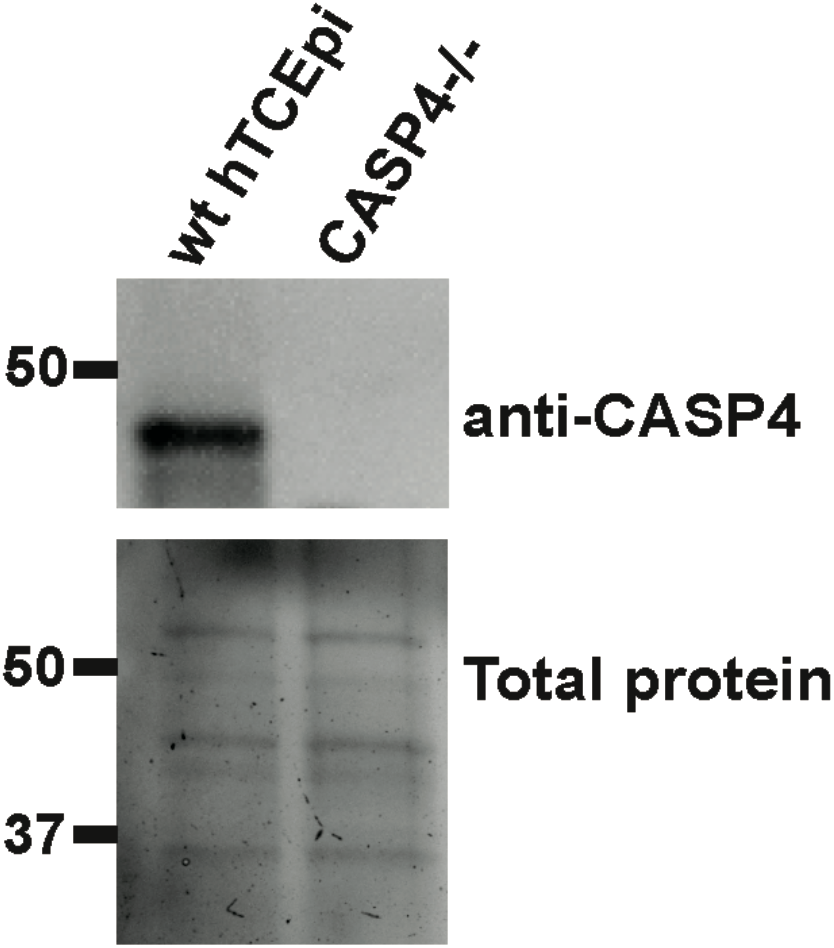
Confirmation of caspase-4 deletion. Lysates from wt hTCEpi cells or caspase-4 candidate knockout cells were collected in RIPA buffer and clarified. Total protein was visualized using Bio-Rad stain free gels prior to transfer. Blot was probed with anti-caspase-4 antibody.

**Supplemental Figure 3.**
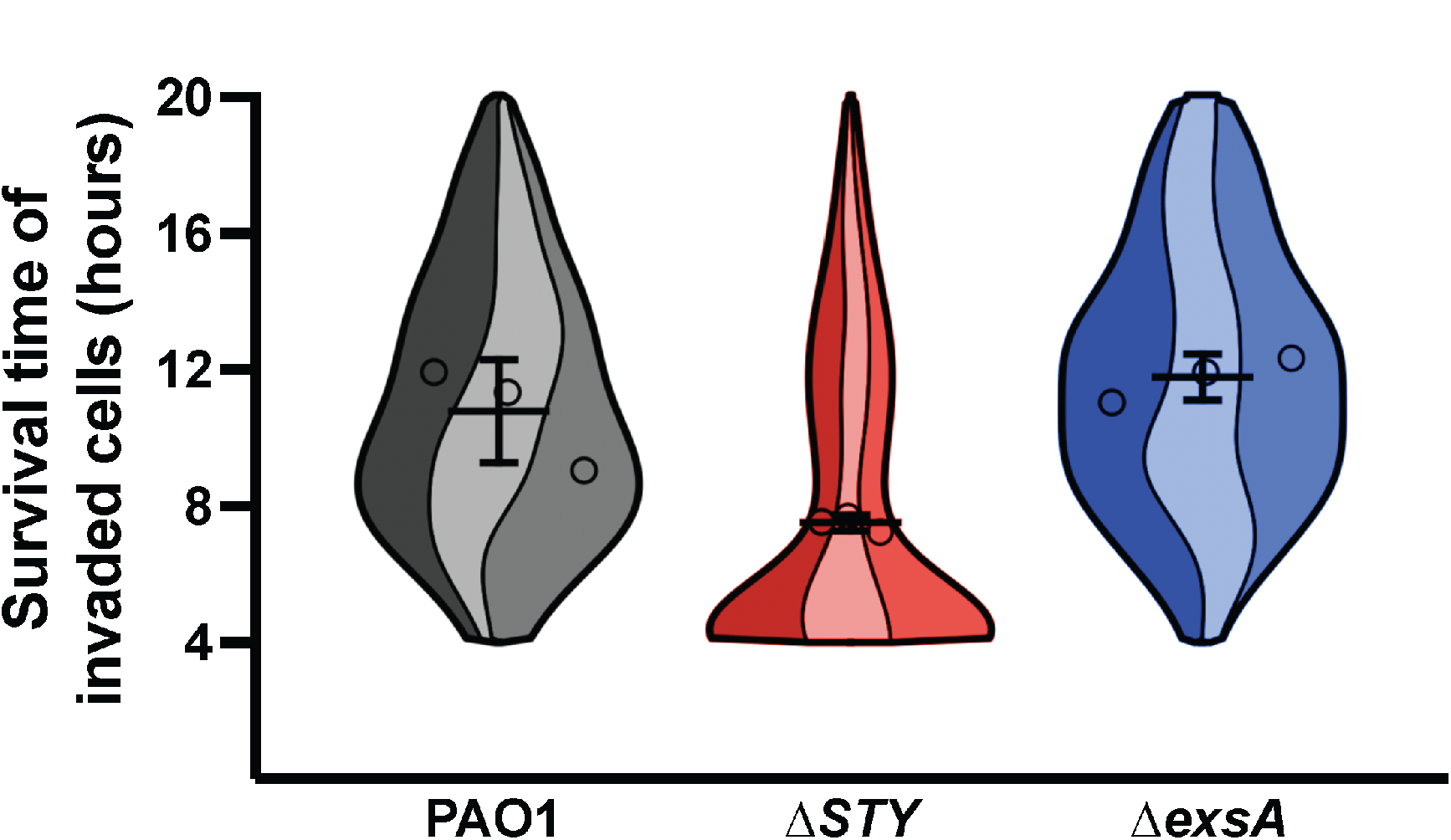
Control monoclonal cells for CRISPR-Cas9 mutagenesis. A control cell line was generated using non-targeting guide RNA, and monoclonal line generated. Cells were infected with wt PAO1, PAO1Δ*exoSTY* (each using the T3SS-GFP reporter) or PAO1Δ*exsA* (pBAD-GFP induced) at an MOI of 10. Nuclei were labeled with Hoechst and Propidium iodide. Extracellular bacteria were eliminated with amikacin at 3 hours post infection, and imaged hourly from 4 to 20 hours post infection. Images were analyzed with a computational approach to segregate invaded cells and measure survival times in hours. Three replicates were combined into a single super violin plot. Mean survival times with SD error bars are displayed.

**Supplemental Figure 4.**
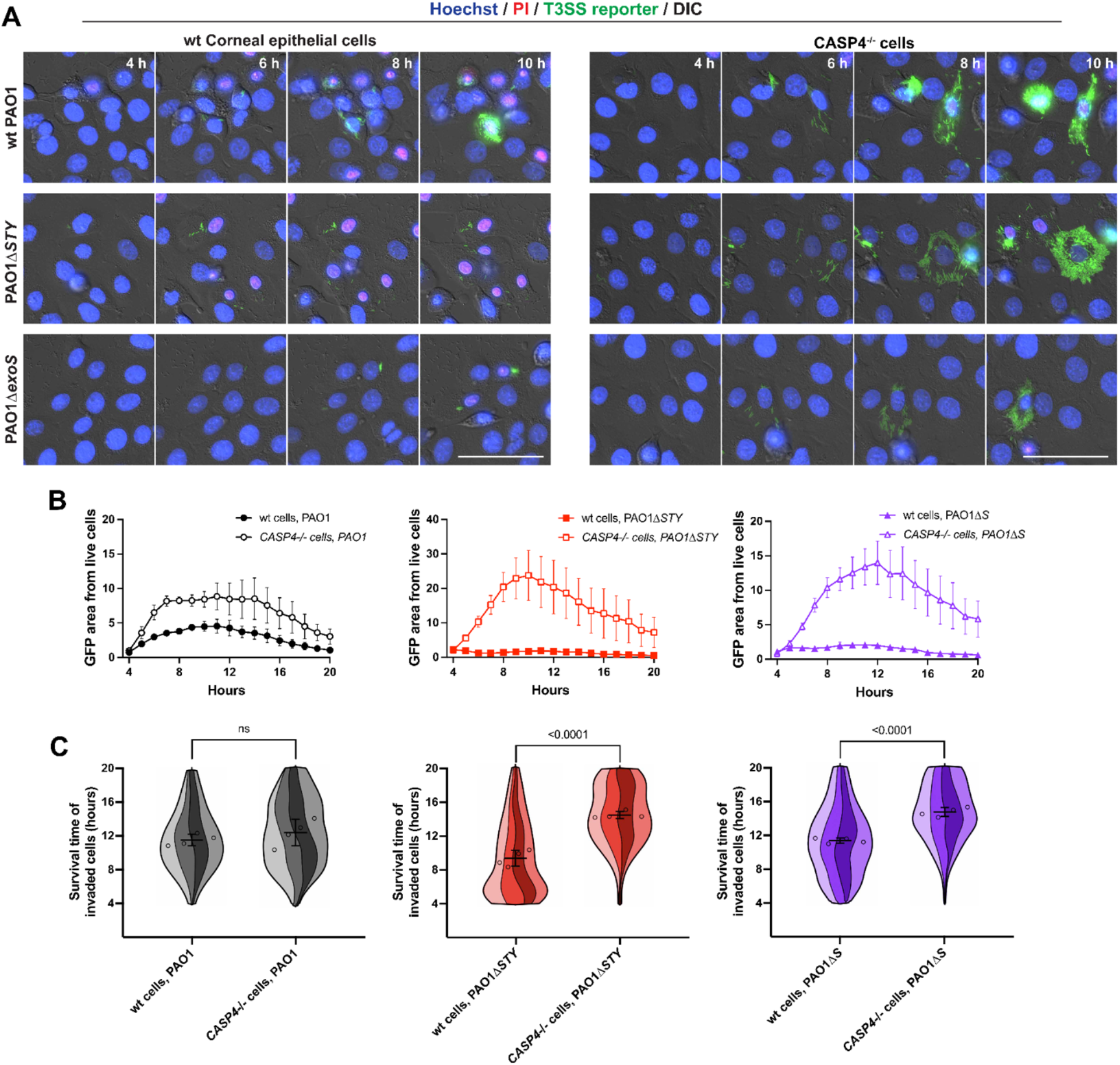
Host cell survival time and intracellular replication of single exotoxin deletion of ExoS. (A) Corneal epithelial cells or cells lacking caspase-4 were infected with wt PAO1, PAO1Δ*exoSTY*, or PAO1Δ*exoS* (each visualized by T3SS-GFP reporter) at an MOI of 10. Nuclei were labeled with Hoechst and propidium iodide. Extracellular bacteria were eliminated with amikacin at 3 hours post infection, and imaged hourly from 4 to 20 hours post infection. Images from 4-10 hours are shown. Scale bar = 50 µm. (B) GFP area was summed within the boundaries of live cells and normalized to PAO1 (wt) at 4 hours post infection. (C) Cells from indicated infections were analyzed with a computational approach to segregate only invaded cells and measure survival times in hours. Mean survival times with SD error bars are displayed, and significance determined by Student’s t-test.

